# Immp2l knockdown increases stimulus-driven instrumental behaviour but does not alter goal-directed learning or neuron density in cortico-striatal circuits in a mouse model of Tourette syndrome and autism spectrum disorder

**DOI:** 10.1101/2023.02.12.528225

**Authors:** Beatrice K. Leung, Sam Merlin, Adam K. Walker, Adam J. Lawther, George Paxinos, Valsamma Eapen, Raymond Clarke, Bernard W. Balleine, Teri M. Furlong

## Abstract

Cortico-striatal neurocircuits mediate goal-directed and habitual actions which are necessary for adaptive behaviour. It has recently been proposed that some of the core symptoms of autism spectrum disorder (ASD) and Gilles de la Tourette syndrome (GTS), such as tics and repetitive behaviours, may emerge because of imbalances in these neurocircuits. We have recently developed a model of ASD and GTS by knocking down *Immp2l*, a mitochondrial gene frequently associated with these disorders. The current study sought to determine whether *Immp2l* knockdown (KD) in male mice alters flexible, goal- or cue-driven behaviour using procedures specifically designed to examine response-outcome and stimulus-response associations, which underlie goal-directed and habitual behaviour, respectively. Whether *Immp2l* KD alters neuron density in cortico-striatal neurocircuits known to regulate these behaviours was also examined. *Immp2l* KD mice and wild type-like mice (WT) were trained on Pavlovian and instrumental learning procedures where auditory cues predicted food delivery and lever-press responses earned a food outcome. It was demonstrated that goal-directed learning was not changed for *Immp2l* KD mice compared to WT mice, as lever-press responses were sensitive to changes in the value of the food outcome, and to contingency reversal and degradation. There was also no difference in the capacity of KD mice to form habitual behaviours compared to WT mice following extending training of the instrumental action. However, *Immp2l* KD mice were more responsive to auditory stimuli paired with food as indicated by a non-specific increase in lever response rates during Pavlovian-to-instrumental transfer. Finally, there were no alterations to neuron density in striatum or any prefrontal cortex or limbic brain structures examined. Thus, the current study suggests that Immp2l is not necessary for learned maladaptive goal or stimulus driven behaviours in ASD or GTS, but that it may contribute to increased capacity for external stimuli to drive behaviour. Alterations to stimulus-driven behaviour could potentially influence the expression of tics and repetitive behaviours, suggesting that genetic alterations to *Immp2l* may contribute to these core symptoms in ASD and GTS. Given that this is the first application of this battery of instrumental learning procedures to a mouse model of ASD or GTS, it is an important initial step in determining the contribution of known risk-genes to goal-directed versus habitual behaviours, which should be more broadly applied to other rodent models of ASD and GTS in the future.

## Introduction

Immp2l (inner mitochondria protein peptidase 2 subunit) is an enzyme encoded by the *Immp2l* gene on chromosome 7 at a location (7q AUTS1) known for its susceptibility to autism spectrum disorder (ASD)(IMGSAC, 1998; Maestrini et al., 2010). As part of the inner membrane peptidase complex, its role is to cleave signal proteins for maturation within the mitochondrial inter-membrane space (Clarke, Furlong, & Eapen, 2020; Clarke, Lee, & Eapen, 2012; Kunova et al., 2022) and it is expressed in most tissues of the body, including the brain (Bertelsen et al., 2014; Gokoolparsadh et al., 2017; Petek et al., 2001). Genetic alteration to *Immp2l* is considered to be a potential risk factor for several neurodevelopmental disorders (Clarke et al., 2020; Kunova et al., 2022). In particular, Immp2l has often been linked to ASD (Casey et al., 2012; Leblond et al., 2019; Maestrini et al., 2010) and Gilles de la Tourette syndrome (GTS) (Bertelsen et al., 2014; Patel et al., 2011; Petek et al., 2001), and to a lesser extent to schizophrenia (Goes et al., 2015) and attention-deficit hyperactivity disorder (ADHD)(Elia et al., 2010). These neurodevelopmental disorders are highly heritable with complex aetiology that is not yet understood, but are often co-morbid with each other, as well as with obsessive-compulsive disorder (OCD), anxiety, and learning disabilities suggesting a shared background (Burd, Li, Kerbeshian, Klug, & Freeman, 2009; Caamano et al., 2013; Freeman et al., 2000; Giovinazzo, Marciano, Giana, Curatolo, & Porfirio, 2013; Kadesjo & Gillberg, 2000; Larson et al., 2010; Mattila et al., 2010; Ueda & Black, 2021).

The defining characteristics of ASD are impaired social interaction and communication, repetitive behaviours, restricted interests, rigid routines and resistance to change, and GTS is defined by vocal and motor tics (e.g., meaningless sounds or words, eye blinking, body contortions, or clapping) (APA, 2013; Rizzo, Gulisano, Domini, C., & Curatolo, 2017; Robertson et al., 2017). There is also overlapping symptomatology between ASD and GTS including the occurrence of motor stereotypies, which are purposeless, repetitive behaviours (such as head nodding or arm flapping which differ to tics), sensory hypersensitivity, and ritualistic, compulsive behaviours associated with OCD (Barry, Baird, Lascelles, Bunton, & Hedderly, 2011; Rizzo et al., 2017; Ubni, Achinivu, Seri, & Cavanna, 2020). The striatum is thought to play a central role in the symptomology of ASD and GTS (Fuccillo, 2016; Lewis, Tanimura, Lee, & Bodfish, 2007; Robertson et al., 2017). For example, the striatum is enlarged in ASD and its size correlates with the severity of repetitive behaviours (Langen et al., 2009), and in GTS the size of the striatum correlates with tic and OCD severity (Bloch, Leckman, Zhu, & Peterson, 2005). Behavioural assessment of our recently developed *Immp2l* knockdown (KD) mouse (Fang, Eapen, & Clarke, 2017) revealed no effect on social interactions, and no effect on marble burying and self-grooming behaviour in male mice, which measure lower-order repetitive behaviour (Kreilaus, Chesworth, Eapen, Clarke, & Karl, 2019; Lawther et al., 2022). However, *Immp2l* KD mice did demonstrate increased locomotor responses to d-amphetamine (Kreilaus et al., 2019; Lawther et al., 2022) which is a striatum-dependent behaviour (Parkinson, Olmstead, Burns, Robbins, & Everitt, 1999; Staton & Solomon, 1984) and hence whether immp2l mediates other striatal-dependent behaviours should be investigated.

The striatum consists of three sub-regions, the caudate, putamen and nucleus accumbens, which form independent circuits with cortical and subcortical limbic structures (Balleine & O’Doherty, 2010; Simmler & Ozawa, 2019; Voorn, Vanderschuren, Groenewegen, Robbins, & Pennartz, 2004). These cortico-striatal limbic circuits are well recognised to play different roles in goal-directed versus habitual behaviour, which are probed using instrumental learning procedures that test sensitivity to outcome value (Balleine & O’Doherty, 2010; Lerner, 2020; Ostlund & Balleine, 2008b; Simmler & Ozawa, 2019). Specifically, rodents are initially trained to press a lever for a food outcome, such as a pellet, and then tested for lever response rates after the value of the outcome is changed, for example by feeding the pellet to satiety (Balleine & O’Doherty, 2010; Perez & Dickinson, 2020; Simmler & Ozawa, 2019). When animals have appropriately learned this response-outcome relationship, responses on the lever associated with the devalued outcome will be reduced compared to when the lever is associated with a food outcome that is still valued, and thus goal-directed behaviour is demonstrated (Balleine & O’Doherty, 2010; Perez & Dickinson, 2020; Simmler & Ozawa, 2019). This procedure has been used to establish the neurocircuitry that mediates goal-directed behaviour in rodents (Balleine & O’Doherty, 2010; Simmler & Ozawa, 2019). It has been shown that lesions or inactivation of the dorsomedial striatum (equivalent to caudate) or the nucleus accumbens, or their associated circuitry, including prelimbic, insular, and orbital cortexes, and basolateral amygdala, will reduce goal-directed behaviour as indicated by a failure to reduce responding on the lever under devalued conditions (Bradfield, Dezfouli, van Holstein, Chieng, & Balleine, 2015; Corbit & Balleine, 2005; Corbit, Leung, & Balleine, 2013; Hart, Bradfield, & Balleine, 2018; Ostlund & Balleine, 2008a; Parkes, Bradfield, & Balleine, 2015).

In contrast, habitual behaviour in rodents is reduced by lesions of the dorsolateral striatum (equivalent to putamen) or its associated circuitry, including infralimbic cortex, dopaminergic nigral neurons, central amygdala and lateral hypothalamus (Becchi, Buson, & Balleine, 2021; Bingul, Merlin, Carrive, Killcross, & Furlong, 2022; Coutureau & Killcross, 2003; Lingawi & Balleine, 2012; Marquez et al., 2020; Yin, Knowlton, & Balleine, 2004). Habitual actions are normally adaptive and occur when an action is well-learned so that it becomes automated and easier to perform with repeated practice without cognitive effort (Lerner, 2020; Robbins & Costa, 2017; Simmler & Ozawa, 2019). Thus, well trained instrumental actions in rodents will become less flexible and stimulus driven rather than outcome driven (i.e., mediated by stimulus-response associations), and will therefore be insensitive to changes in outcome value when tested following outcome devaluation in contrast to goal-directed behaviour (Adams, 1982; Dickinson, Balleine, Watt, Gonzalez, & Boakes, 1995). Of particular interest, performance on translational versions of these tasks are altered in individuals with ASD, GTS, OCD, which is suggestive of impaired goal-directed learning due to altered sensitivity to devaluation, or capacity for learning response-outcome or stimulus-response relationships (Alvares, Balleine, Whittle, & Guastella, 2016; Delorme et al., 2016; Gillan et al., 2011; Scholl, Baladron, Vitay, & Hamker, 2022). Further, it has been proposed that tics may resemble habitual behaviour as they are automated, somewhat intentional actions that occur in response to sensory stimuli, and thus may share common neural mechanisms (Delorme et al., 2016; Scholl et al., 2022; Singer, 2013). An imbalance between goal-directed and habit behaviour has also been proposed to underlie inflexible, compulsive and repetitive behaviour, which may be triggered by external stimuli rather than intended goals (Alvares et al., 2016; Gillan et al., 2011; Hadjas, Luscher, & Simmler, 2019).

The aim of the current study was to determine whether *Immp2l* KD alters cortico-striatal circuits in male mice, given the higher prevalence of ASD and GTS in male than female individuals (Burd et al., 2009; Freeman et al., 2000; Kadesjo & Gillberg, 2000). Goal-directed behaviour and cognitive flexibility were first examined following instrumental training by testing for sensitivity to outcome value, and by altering the response-outcome relationship through reversal learning and contingency degradation (Bradfield, Bertran-Gonzalez, Chieng, & Balleine, 2013; Furlong et al., 2016; Ostlund & Balleine, 2008b). Pavlovian-to-Instrumental Transfer task (PIT) was also used to examine the impact of environmental stimuli on goal-directed behaviour (Holmes, Marchand, & Coutureau, 2010; Laurent, Leung, Maidment, & Balleine, 2012; Leung & Balleine, 2015; Ostlund & Balleine, 2008b). The capacity to form habitual actions was then examined by testing sensitivity to outcome value following extended instrumental training of separate animals (Dickinson et al., 1995; Hadjas et al., 2019). Finally, neuron density in cortico-striatal limbic circuits was examined to determine whether *Immp2l* KD recapitulates neuroanatomical changes in ASD or GTS, which includes reduced neuron density in the striatum and increased neuron density in the prefrontal cortex (Courchesne et al., 2011; Falcone et al., 2021; Kalanithi et al., 2005; Kataoka et al., 2010; Wegiel et al., 2014). This is the first application of goal-directed and habitual tests of behaviour to a genetic animal model of ASD or GTS as operationalised by Dickinson, Balleine and colleagues (Dickinson & Balleine, 1994; Dickinson et al., 1995; Ostlund & Balleine, 2008b; Perez & Dickinson, 2020), which is an important step toward probing translational striatal-dependent alterations in behaviour in these mouse models.

## Materials and Methods

### Overview of experiments

Two experiments were conducted to assess the effects of *Immp2l* knockdown (KD) on cognitive flexibility and goal-directed behaviour in male adult mice. In Experiment 1, the behavioural assessments were conducted using two levers and ratio schedules of reinforcement that promote goal-directed actions (Perez & Dickinson, 2020). In Experiment 2, for a separate cohort of mice, the behavioural assessments were conducted using a single lever and an interval schedule of reinforcement to promote habitual behaviour (Hadjas et al., 2019; Perez & Dickinson, 2020). A timeline of the experiments is shown in Figure 1. In Experiment 3, a separate cohort of male adult mice were used to examine neuron density in cortico-striatal limbic circuits using immunohistochemistry.

**Figure 1:**
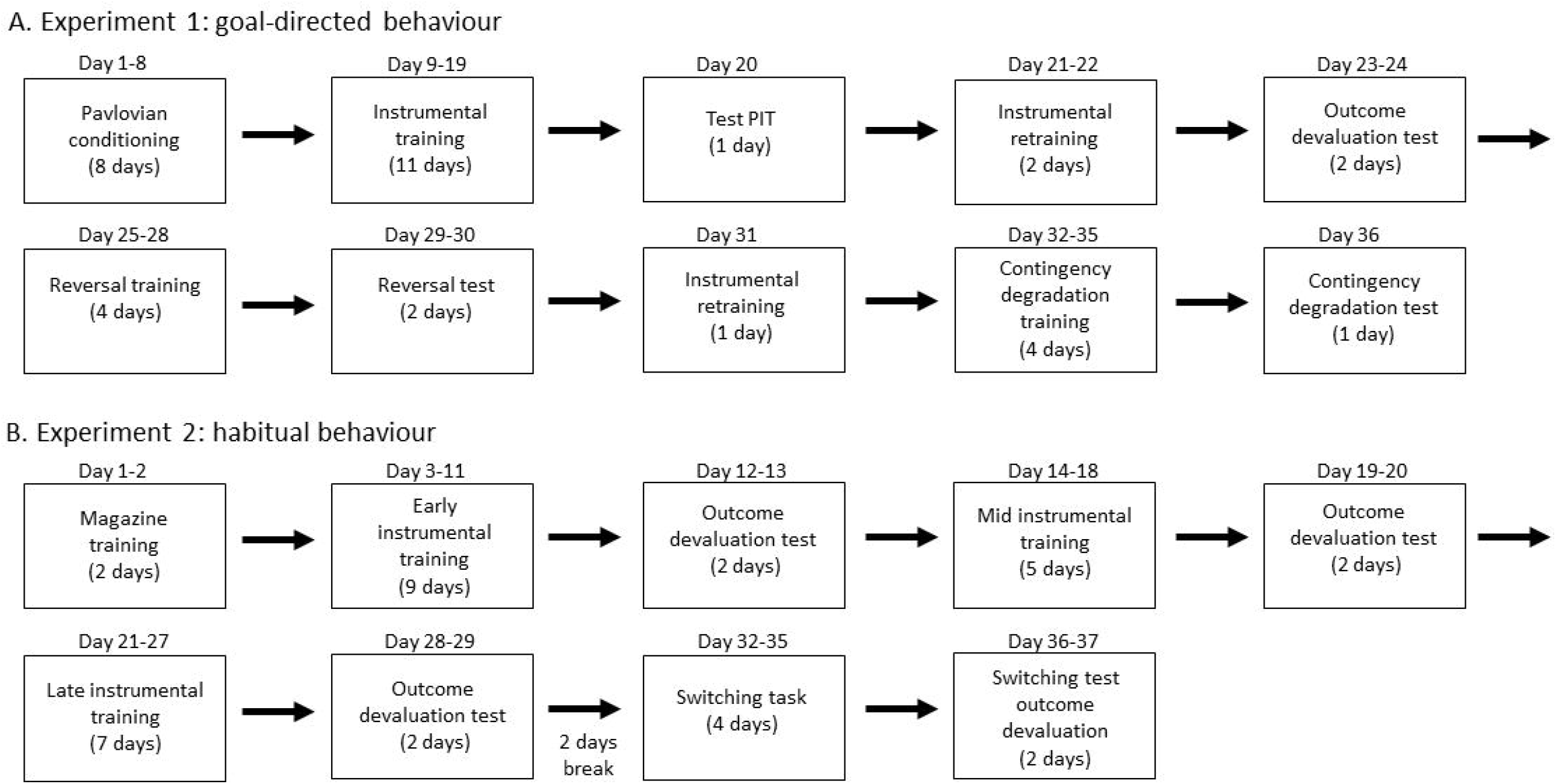
Timeline of experiments. a) Timing of behavioural procedures used to examine goal-directed behaviour which consisted of Pavlovian conditioning and instrumental training with two leveres, and tests for specific PIT, outcome devaluation, reversal learning and contingency degradation. b) Timing of behavioural procedures used to examine habitual behaviour. This includes three outcome devaluation tests following varied lengths of instrumental training on a single lever, and a switching task in which a new lever was introduced. The number of days since commencement of the experiment is indicated above each box, and the number of days dedicated to each stage of the experiment is indicated within the box. See methods for details of each behavioural procedure.

### Subjects

Male *Immp2l* KD mice (KD; Experiment 1 (n=11), Experiment 2 (n=14), and Experiment 3 (n=14)) and wild type-like mice (WT; Experiment 1 (n=11), Experiment 2 (n=14), Experiment 3 (n=18)) were used in this study (aged 4-5 months old). Mice were bred and housed at the Ingham Institute Biological Resource Unit (Liverpool, Australia) until they were adults and then transported to the animal facility at University of New South Wales (Kensington, Australia; Experiments 1 and 2) or at Neuroscience Research Australia (Randwick, Australia; Experiment 3). Mice were group housed (2–4 per cage) in individually ventilated cages (GM500 Green, Techniplast) with corn cob bedding and enrichment, including crinkle cut cardboard nesting material, a red igloo and a blue tunnel (Bioserv Biotechnologies) and kept on a 12 h light dark cycle (lights on between 7:00 A.M. and 7:00 P.M.). All procedures were approved by the University of New South Wales Ethics Committee (ethics #19/81A), in accordance with the Australian Code of Practice for the Care and Use of Animals for Scientific Purposes.

### Behavioural apparatus

For all behavioural experiments, training was conducted in 24 operant chambers enclosed in sound- and light-attenuating cabinets (MED Associates, USA). Each chamber was fitted with a pellet dispenser capable of delivering a 20 mg grain food pellet (F0163, BioServ Biotechnologies), and a pump fitted with a syringe outside the chamber capable of delivering 0.025 mL 20 % sucrose solution (white sugar, Coles Supermarket; dissolved in H_2_O) into a recessed magazine inside the chamber. The chambers were fitted with two retractable levers that could be inserted individually on the left and right sides of the magazine. Head entries into the magazine were detected via an infrared photobeam. The chambers also contained a sonalert 3 kHz tone generator and a white noise amplifier to provide stimuli for Pavlovian conditioning. Operant chambers were fully illuminated during all experimental stages by a 3 W, 24 V house light located on the upper edge of the wall opposite to the magazine except during the delivery of the sucrose during which the house light was turned off. All training sessions were pre-programmed on computers located in a separate room through MED Associates software (Med-PC). These computers also recorded the experimental data from each session.

### Food restriction

Mice in Experiment 1 and 2 underwent 5 days of food restriction prior to the onset of behavioural training, and this continued throughout the duration of the experiment. During this time, they received 0.7-2 g of standard chow daily until the end of the experiment in order to maintain body weight at approximately 85% of their pre-restriction body weight. Mice utilized in Experiment 3 were not food deprived.

## Experiment 1: goal-directed behaviour

### Pavlovian training

Mice received 8 daily sessions of Pavlovian training (Furlong et al., 2016; Laurent et al., 2012; Leung & Balleine, 2015). Each session was 1 hour duration and consisted of presentations of the two conditioned stimuli (CS; i.e., a tone or a white noise), each paired with the delivery of either the sucrose or the pellet outcomes. Each CS lasted 2 minutes and was presented four times in a pseudorandom order with a variable intertrial interval of 5 minutes. Half of the mice in each group received tone–pellet and noise–sucrose pairings, whereas the other half received the opposite CS–outcome pairings. The appropriate outcome was delivered during the CS on a random-time 30-second schedule.

### Instrumental training for goal directed behaviour

Mice next received 11 days of instrumental training during where two actions (left and right lever press responses) were trained with the different outcomes (i.e., pellets or sucrose) in two separate daily sessions (Furlong et al., 2016; Laurent et al., 2012; Leung & Balleine, 2015). Half of the mice in each group received left lever press and pellets in one session and right lever press and sucrose in the other session, whereas the remainder received the opposite action–outcome relationships. The order of the training sessions was varied over days. Each session ended when 20 outcomes were earned or when 30 minutes had elapsed. For the first 2 days, lever pressing was continuously reinforced. Thereafter, the probability of the outcome given a response [p(O/R)] was gradually shifted over days using an increasing random ratio (RR) schedule: a RR5 schedule (p0.2) was used on days 3–5, a RR10 (p0.1) schedule was used on days 6–8, and a RR20 (p0.05) schedule used on days 9–11.

### Pavlovian-instrumental transfer test

After the final day of RR20 training, mice were given a Pavlovian-instrumental transfer (PIT) test (Furlong et al., 2016; Laurent et al., 2012; Leung & Balleine, 2015). Both levers were inserted into the chamber, but no outcomes were delivered during the test. Responding was extinguished on both levers for 10 minutes to reduce the baseline rate of performance after which each CS was presented four times over the next 40 minutes in the following order: tone–noise–noise–tone–noise–tone–tone–noise. Stimulus presentations lasted 2 minutes and were separated by a 3-minute fixed intertrial interval. Specific PIT is demonstrated when lever responses are increased during the CS specifically on the lever that is associated with the same outcome predicted by the CS (Laurent et al., 2012).

### Outcome devaluation tests

Mice were then retrained on the RR20 schedule for 2 consecutive days. The following day, they received ad libitum access to one of the two outcomes, either pellets or sucrose, for 1 hour in distinct feeding cages located in a room different from where training had been conducted (Furlong et al., 2016; Laurent et al., 2012). Half of the mice in each action–outcome assignment received pellets (5 g placed in a dish), and the remaining mice received sucrose (100 ml in a drinking bottle). The mice were then given a 5-minute choice extinction test in which both levers were available, but no outcomes were delivered. The same procedure was repeated 1 day later except that the mice were given ad libitum access to the other food outcome so that all mice were tested under devalued and valued conditions in counter-balanced order. Goal-directed behaviour is demonstrated when animals reduce responding under devalued conditions compared to valued conditions (Furlong et al., 2016; Laurent et al., 2012).

### Instrumental reversal learning

Following outcome devaluation, mice received 4 days of instrumental training on a RR20 schedule where the response-outcome relationships were reversed (Bradfield & Balleine, 2017; Furlong et al., 2016). That is, mice that had the left lever paired with pellets and right lever paired with sucrose, now had the left lever paired with sucrose and the right lever paired with pellets, and vice versa. After training, mice underwent the same outcome devaluation tests as described above. Animals that are flexible in their behaviour are expected to demonstrate outcome devaluation that is consistent with the new/reversed response-outcome contingency rather than the initially trained contingency (Furlong et al., 2016).

### Contingency degradation

Finally, mice were given one day of RR20 retraining before 4 days of contingency degradation training. Similar to instrumental training sessions, mice received two separate daily sessions with the response-outcome contingencies (Bradfield et al., 2013; Furlong et al., 2016). However, during these sessions one of the two outcomes was also freely delivered at the same probability as a lever press (p0.05). Half of the animals with each response-outcome pairings had the pellet freely delivered and half had the sucrose freely delivered in order to degrade the response-outcome contingency. After the final day of degradation training, mice were given a 5-minute choice test in extinction where both levers were available, but no outcomes were delivered. Animals that are flexible in their behaviour are expected to be sensitive to contingency degradation and to reduce responding on the degraded lever compared to the lever where the response-outcome contingency was not degraded (Bradfield et al., 2013; Furlong et al., 2016).

### Data analyses

All data were analysed using mixed-model ANOVAs followed by simple main effects analyses to establish the source of any significant interactions. For Pavlovian conditioning, magazine entries for each session were analysed as an elevation ratio, calculated as magazine entries during CS presentations divided by total number of magazine entries during the CS periods and the Pre CS periods of equal duration. For instrumental training, lever press responding was analysed as rate of pressing each day after responses on the two levers were averaged for each day. For the PIT test, data was analysed as raw lever responses during CS presentations, and as Pre CS periods (of equal duration) as a measure of baseline responding. For all other behavioural tests, data was analysed as lever responses at test divided by baseline responding. For outcome devaluation tests, the baseline responding was calculated using the average rate of responding on the last two days of training prior to the test. For contingency degradation test the baseline responding was the rate of responding from the 1 day of retraining.

## Experiment 2: habitual behaviour

### Magazine training

A separate group of mice to those tested for goal-directed behaviour in Experiment 1 were first given 2 sessions of magazine training where 20 pellets were delivered on a random time 60-second schedule.

### Instrumental training for habitual behaviour

Mice received extended instrumental training for a total of 21 days during which one action (left or right lever press responses) earned a pellet outcome. Half of the mice in each group received the left lever and half received the right lever. Sessions ended after 25 outcomes were earned or 80 minutes had elapsed. For the first 3 days, lever pressing was continuously reinforced. Thereafter, the rate of reinforcement shifted to a random interval (RI) schedule where outcomes were delivered for the first lever press made after an average interval of 15 (RI15), 30 (RI30) or 60 (RI60) seconds in order to promote habitual behaviour (Bingul et al., 2022; Hadjas et al., 2019). Mice received 2 days of RI15, 4 days of RI30 and 12 days of RI60 training.

### Outcome devaluation tests

Mice were given three sets of outcome devaluation tests throughout instrumental training to assess the development of habitual behaviour. Tests were given after the last day of RI30 (early training), after day 5 of RI60 (mid training) and after day 12 of RI60 (late training). During outcome devaluation, half of the mice were given ad libitum access to pellets (5 g) and the other half received home chow (5 g) for 1 hour in distinct feeding cages located in a room different from where training had been conducted (Hadjas et al., 2019; Mo et al., 2020). The mice were then given a 5-minute extinction test where the lever was presented but no outcomes were delivered. The same procedure was repeated 1 day later except that mice were given ad libitum access to the other outcome so that each mouse was tested under devalued and valued conditions in counter-balanced order (Mo et al., 2020). Mice that are habitual in their responding will press the lever equally under devalued and valued conditions (Hadjas et al., 2019; Mo et al., 2020).

### Switching task

Following the last devaluation test, mice received 4 days of instrumental training where both the left and right levers were presented. During this switching task, the originally trained lever became inactive and no longer earned outcomes, and the new lever became the active lever and now earned outcomes on a RI60 schedule. Following 4 days of training, animals were tested following outcome devaluation. This test was identical to the outcome devaluation tests (above) except that both levers were presented. It was expected that if animals were habitual in their responding they would be slower to switch their preference from the original/inactive lever to the new/active lever during training as occurs during reversal of response outcome-contingencies (Gourley, Lee, Howell, Pittenger, & Taylor, 2010).

### Data Analyses

All data were analysed using mixed-model ANOVAs followed by simple main effects analyses to establish the source of any significant interactions, as described for Experiment 1.

## Experiment 3: neuron density

### Histology

Naïve mice were utilised for examination of neuron numbers using NeuN (neuronal nuclear protein) in cortico-striatal limbic circuits known to regulate goal-directed and habitual behaviour. Mice were euthanised with an over-dose of pentobarbitone (20 mg) and transcardially perfused with 0.9 % sodium chloride followed by 40 ml of 4 % paraformaldehyde (0.1 M phosphate buffer, pH=7.4; PB). The brains were excised and post-fixed in paraformaldehyde for 1 hr followed by 20 % sucrose (in PB) for cryoprotection for 2 days. The brains were then sectioned on a freezing cryostat (Leica Biosystems, 35 μm thick) with sections collected every 105 μm for each brain structure analysed.

For immunohistochemistry (Sheth, Furlong, Keefe, & Taha, 2016), the sections were processed in 50 % ethanol (30 min), 3 % H_2_O_2_ in 50 % ethanol (30 min) and then 5 % normal horse serum (NHS, 30 min) before incubating in primary antibody (mouse-anti-NeuN, Merck Millipore, #MAB377; 1:5000 for 24 hrs in 5 % NHS, 2 % triton-X in PB). Sections were then washed in PB (3x 20 min) before incubation in secondary antibody (biotinylated donkey-anti-mouse, Jackson Immunoresearch, #715065150; 1:1000 for 24 hrs) and then Vectastain ABC reagent (Vector Labs; 1:750 for 2 hrs) (both in 5 % NHS, 2 % triton-X in PB). Finally, a DAB (diaminobenzidine) reaction was used to visualise a black reaction product where sections were placed in a 0.1 M sodium acetate buffered solution (pH=6.0) containing 0.5 % DAB, 0.04 % ammonium chloride, 1 % nickel sulphate, 0.2 % D-glucose and 735 units/ml of glucose oxidase (for 10 min). Sections were then mounted on gelatine-coated slides and coverslipped with Entellan.

### Quantification

Slides were scanned using an Aperio Scanscope slide scanner (20X objective; Leica Biosystems). NeuN-positive cells were quantified using the thresholding tool in Image J software (National Institutes of Health) by an experimenter blind to group. The same threshold was used for all animals for each brain region quantified. Regions were quantified bilaterally at two rostro-caudal levels according to a mouse brain atlas (Franklin & Paxinos, 2008). For the PFC, the cingulate, prelimbic, infralimbic, lateral orbital and anterior insular cortices were quantified between bregma 1.8 mm and 2.00 mm. For the striatum, DLS and NA-shell and NA-core were quantified between 1.2 mm and 1.4 mm, and the DMS between 0.4 mm and 0.6 mm. The ventral pallidum was quantified between 0.1 mm and 0.3 mm, the lateral hypothalamus and basolateral and central amygdala between 1.4 mm and -1.6 mm, and the ventral tegmental area and substantia nigra compacta between -3.3 mm and -3.1 mm.

### Data Analyses

Total NeuN for each animal was determined by averaging the counts across the sections examined for each brain region which were expressed per mm^2^ of area examined. Individual one-way ANOVAs were used to determine whether there were differences between KD and WT groups for each brain region examined.

## Results

### Experiment 1: goal-directed behaviour

#### Pavlovian conditioning

Pavlovian conditioning was assessed using head entries to the magazine during CS presentations compared to the pre-CS period and are shown as elevation ratios in Figure 2a. It was found that both WT and KD mice increased their head entries to the magazine during the CS period as training progressed (F _(1, 20)_ = 23.379; *p* < 0.001) and there was no main effect of genotype (F _(1, 20)_ = 0.066; *p* = 0.80), but there was a session x genotype interaction (F _(1, 20)_ = 2.290; *p* = 0.031). Simple main effect analysis on each training session revealed a significant difference between groups only on session 3 (F _(1, 20)_ = 4.899; *p* = 0.039).

**Figure 2.**
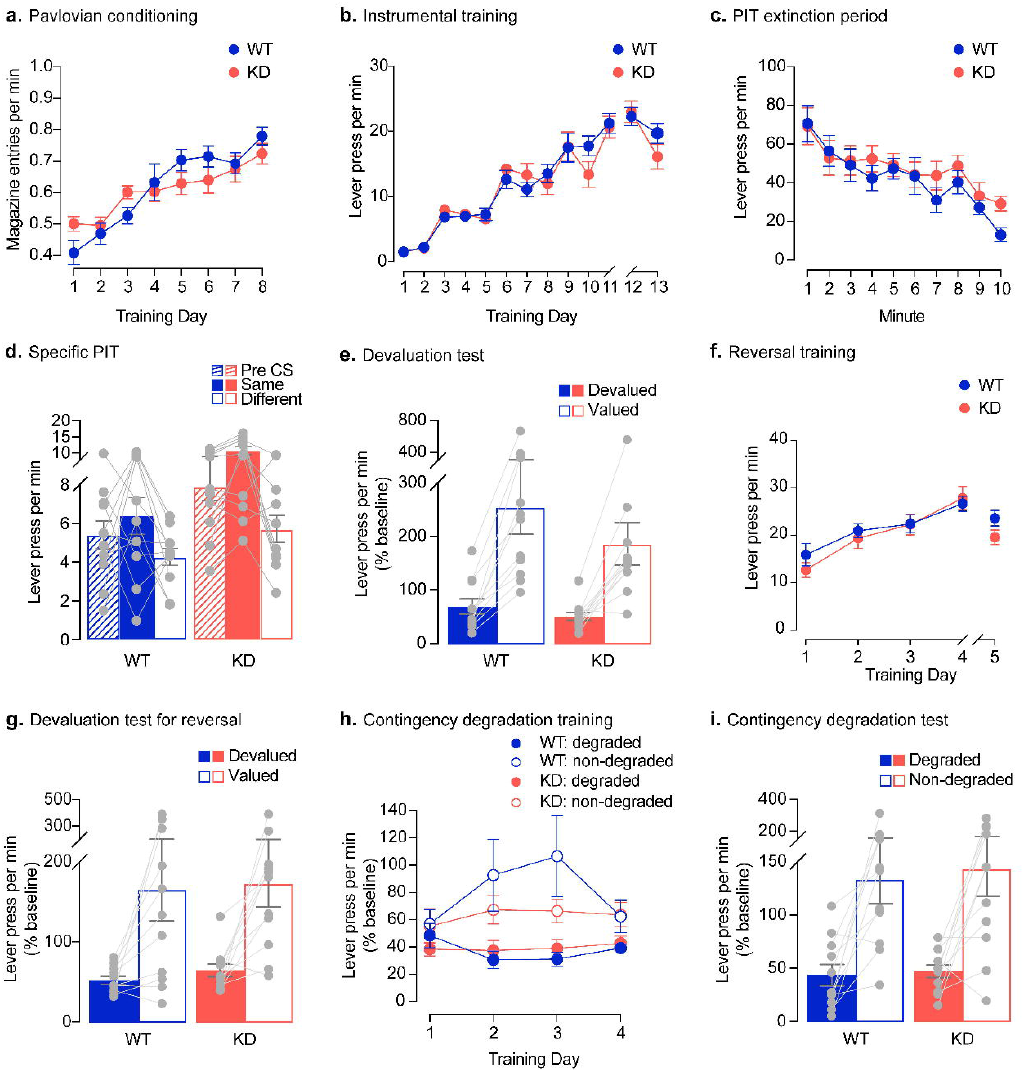
Immp2l knockdown does not impair goal-directed actions. a) Elevation ratio of magazine entries during CS presentations show that both genotypes demonstrated equivalent Pavlovian conditioning across sessions. b) Lever presses across instrumental training and retraining sessions show that both genotypes demonstrated equivalent instrumental conditioning. c) Lever presses during the extinction period of the PIT test show that both genotypes demonstrated equivalent extinction of instrumental responses. d) Lever presses during PIT test separated as the Pre CS period and during the CS period for the lever associated with the same (Same) and different (Different) outcome the CS is predicting. Despite an overall increased lever press rate from KD mice during both the pre-CS and CS periods, both genotypes demonstrated specific PIT, responding more on the lever associated with the same outcome as the CS (Same) than the lever associated with a different outcome to the CS (Different). e) Lever presses during the choice test following outcome devaluation. Both genotypes demonstrated goal-directed behaviour responding more on the lever when it was associated with the still valued outcome (Valued) than when it was associated with the devalued outcome (Devalued). f) Both genotypes demonstrated equivalent lever response rates across reversal training and retraining when the action-outcome contingencies were reversed. g) Lever presses during the choice test following outcome devaluation for reversal learning. Both genotypes demonstrated reversal learning, responding more on the lever when the outcome was still valued (Valued) compared to devalued (Devalued), consistent with the new response-outcome contingency. h) Lever presses during contingency degradation training. Both genotypes demonstrated reduced responding on the lever associated with the outcome that was freely delivered (Degraded) compared to the other lever (Non-degraded). i) Lever presses during the choice test following contingency degradation training. Both genotypes were sensitive to contingency degradation given their greater preference for the lever associated with non-degraded (Non-degraded) outcome compared to the degraded outcome (Degraded). Data are shown as mean ± SEM.

#### Instrumental training

Instrumental training was assessed using the average rate of lever pressing on each lever on each day (Figure 2b). Both genotypes were able to acquire the instrumental response and increase responding on the levers across training sessions (F _(10, 200)_ = 87.023; *p* < 0.001) with no differences between groups (F _(1, 20)_ = 0.02; *p* = 0.89) and no interaction (F _(10, 200)_ = 1.742; *p* = 0.074). Rates of lever pressing was also similar when animals received retraining sessions prior to outcome devaluation (day 12-13 in Figure 2b). There was a decrease in lever press rate on the second day of retraining (F _(1, 20)_ = 14.526; *p* = 0.001) but no differences between genotype (F _(1, 20)_ = 0.540; *p* = 0.471) or interaction (F _(1, 20)_ = 3.022; *p* = 0.097).

#### Pavlovian Instrumental Transfer

Mice were given a PIT test to assess their ability to use stimuli to guide their instrumental actions. To reduce their responding to a low baseline, mice were given 10 minutes of extinction on both levers prior to the onset of the first CS. As shown in Figure 2c, both groups reduced their responding on the levers across the 10-minute extinction period (F _(9,180)_ = 11.72; *p* < 0.001) and there was no main effect of genotype (F _(1, 20)_ = 0.645; *p* = 0.431) or interaction (F _(9, 180)_ = 0.679; *p* = 0.727). During the PIT test, as shown in Figure 2d, we observed a significantly higher baseline rate of responding for the KD group compared to WT group during the pre-CS periods at test (F (1, 21) = 5.952; p = 0.024). This increased responding was also evident during the CS periods, and while both groups were able to demonstrate specific PIT and responded more on the lever associated with the same outcome that the CS predicted compared to the lever that was associated with different outcome (same > different; F (1, 20) = 16.919; p = 0.001), there was a main effect of genotype where the KD responded more than WT overall during the CSs (F (1, 20) = 10.170; p = 0.005) but no interaction between groups (F (1, 20) = 2.597; p = 0.123). Thus, the KD group made more lever responses than the WT group during both the Pre CS and CS periods following equivalent extinction.

#### Outcome Devaluation

Mice were then given an outcome devaluation test to assess their ability to use the value of the outcome to guide their instrumental actions. During the free feeding period prior to the lever choice test, there were no differences between WT and KD mice in the amount of grain pellets (0.95 g vs 0.99 g ; F _(1,20)_ = 0.128; p = 0.724) or sucrose solution (1.81 g vs 2.04 g ; F _(1,20)_ = 2.793; p = 0.110) consumed. During the choice test, the overall lever response rates were similar between WT and KD mice (14.35 vs 15.01 lever presses per min, F _(1, 20)_ = 0.17, p = 0.685). When lever pressing was analysed as responding on the valued versus devalued lever (Fig 2e), it was found that both groups of mice correctly responded on the still valued lever compared to the devalued lever (valued > devalued; F _(1,20)_ = 37.052; *p* < 0.001) and there were no differences between genotype (F _(1, 20)_ = 1.249; *p* = 0.277) or interaction between the two factors (F _(1, 20)_ = 0.915; *p* = 0.350).

#### Reversal learning

Next, we assessed *Immp2l* KD mice on whether they were able to update action-outcome associations by reversing the action-outcome relationships. We found that *Immp2l* KD did not impair reversal learning and animals performed similarly to their WT counterparts. During the four training sessions with the reversed action-outcome contingency, both groups of mice increased their responding on the levers across days (Figure 2f; F _(3, 60)_ = 49.541; *p* < 0.001), but there was no effect of genotype (F (1, 20) = 0.143; *p* = 0.709) or interaction between the two factors (F _(3, 60)_ = 1.525; *p* = 0.217). During the devaluation tests, there were no differences between WT and KD mice in the amount of grain pellets (1.00 g vs 1.15 g ; F _(1,20)_ = 0.823; p = 0.375) or sucrose solution (1.45 g vs 1.56 g ; F _(1,20)_ = 0.620; p = 0.440) consumed, or in the overall lever response rates (15.63 vs 17.92; F = 0.705, p = 0.411). When analysing lever pressing separately, mice were able to correctly respond more on the valued lever compared to the devalued lever based on the new contingencies (Figure 2g; F _(1, 20)_ = 19.629; *p* = 0.001), with no effect of genotype (F _(1, 20)_ = 0.173; *p* = 0.682) or interaction (F _(1, 20)_ = 0.009; *p* = 0.927)

#### Contingency degradation

Finally, we assess the sensitivity of the mice to changes in the contingency between their actions and the outcome by selectively degrading the contingency on one of the two levers. During degradation training, both groups of mice continued to press the lever that delivered the non-degraded outcome, and responded less on the lever where the contingency was degraded by free delivery of the outcome (Figure 2h; non-degraded > degraded; F _(1, 20)_ = 10.708; *p* = 0.004). While there was no effect of training session (F _(3, 60)_ = 1.245; *p* = 0.301), there was a lever by session interaction showing the difference between levers increased over days (F _(3, 60)_ = 3.821; *p* = 0.014). There was no effect of genotype (F _(1, 20)_ = 0.592; *p* = 0.451) or any interactions involving genotype as a factor (All Fs < 1.86; *p* > 0.146). For the contingency degradation test, both groups of mice clearly preferenced their responding to the lever where the response-outcome contingency was intact over the lever where the contingency was degraded (Figure 2i; non-degraded > degraded; F _(1, 20)_ = 25.373; *p* < 0.001), and there was no effect of genotype (F _(1, 20)_ = 0.146; *p* = 0.706) or interaction between the two factors (F _(1, 20)_ = 0.027; *p* = 0.872).

### Experiment 2: habitual behaviour

#### Magazine training

Both groups of mice were able to familiarize themselves to the magazine where outcomes were delivered (Figure 3a). Entries to the magazine increased on session 2 (F _(1, 26)_ = 38.808; *p* < 0.001) with no effect of genotype (F _(1, 26)_ = 2.734; *p* = 0.110) or interaction (F _(1, 26)_ = 0.877; *p* = 0.358).

**Figure 3.**
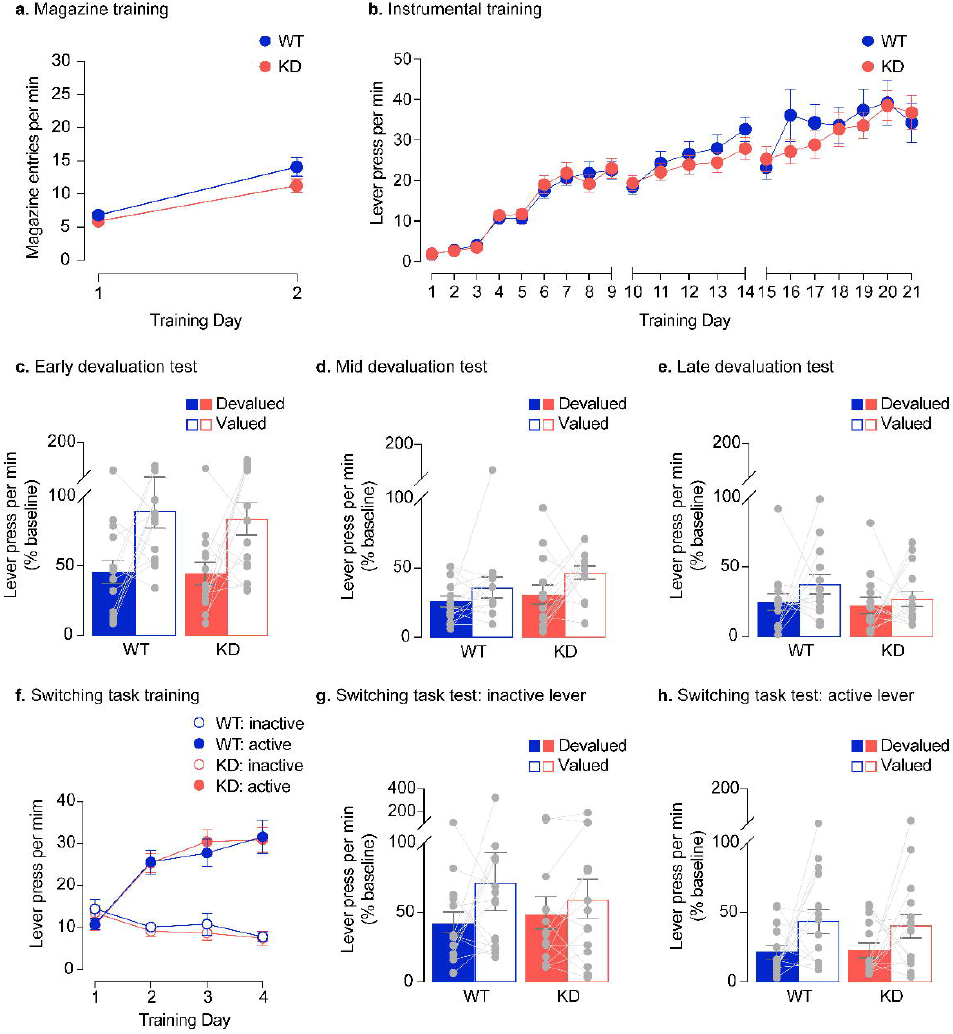
Immp2l knockdown does not impair the development of habitual actions. a) Magazine entries during magazine training did not differ between genotypes. b) Lever presses across early, mid and late instrumental training did not differ between genotypes. c) Lever presses during the test following outcome devaluation after early training. Both genotypes demonstrated goal-directed action, pressing the lever more when the associated outcome was still valued (Valued) versus when the outcome had been devalued (Devalued). d) After mid training, both genotypes also demonstrated goal-directed actions. e) After late training, both genotypes demonstrated habitual actions where lever response rates were equivalent under devalued conditions (Devalued) as they were under valued conditions (Valued). f) Lever presses on the active and inactive levers during training of the switching task. Both genotypes switched their preference to the active lever that delivered an outcome when pressed, over the inactive lever which did not deliver an outcome. g) Lever presses on the inactive lever at test when the outcome was still valued or devalued. For both genotypes lever response rates were equivalent under devalued conditions (Devalued) as they were under valued conditions (Valued), and hence animals were not goal-directed toward the inactive lever. h) Lever presses on the active lever at test when the outcome was still valued or devalued. Both genotypes demonstrated greater responding under valued conditions (Valued) compared to devalued conditions (Devalued) and hence were goal-directed toward the active lever. Data are shown as mean ± SEM.

#### Instrumental training

Both groups of mice also showed similar rates of lever pressing on the RI schedule during training (Figure 3b). Each set of training sessions, between each devaluation test, were analysed separately: sessions 1-9 (early training), sessions 10-14 (mid training) and sessions 15-21 (late training). Throughout training, the two groups of mice performed similarly to each other and increased responding across sessions during early (F _(8, 208)_ = 90.061; *p* < 0.001), mid (F _(4, 104)_ = 16.476; *p* < 0.001) and late (F _(6, 156)_ = 10.150; *p* < 0.001) training. There was no effect of genotype for any training period (All Fs < 0.532; *p* > 0.46) or any interactions with genotype as a factor (All Fs < 2.022; *p* > 0.066).

#### Outcome Devaluation

Mice were given 3 separate devaluation tests following early, mid and late training (Figures 3c, 3d and 3e, respectively). There was no difference in the amount of food consumed between groups in the free-access devaluation period prior to any of these tests (all Fs < 1.215, p > 0.28; data not shown) or in overall lever response rates at test prior (all Fs < 0.772; *p* > 0.388; data not shown). At early training, both groups of mice were able to show appropriate outcome devaluation and responded more on the lever when the outcome was still valued compared to when the outcome was devalued (Figure 3c; valued > devalued; F _(1, 26)_ = 13.271; *p* = 0.001). There was no effect of genotype (F _(1, 26)_ = 0.145; *p* = 0.706) or interaction between the two factors (F _(1,26)_ = 0.037; *p* = 0.849). At mid training, both groups of mice showed a marginal devaluation effect (Figure 3d; valued > devalued; F _(1, 26)_ = 4.587; *p* = 0.042) with no effect of genotype (F _(1, 26)_ = 1.703; *p* = 0.203) or interaction between the two factors (F _(1,26)_ = 0.216; *p* = 0.646). Finally, at late training following extensive RI60 training, both groups of animals were no longer sensitive to changes in the value of the outcome and responded on the lever equally under devalued and valued conditions (Figure 3e, valued = devalued; F _(1, 26)_ = 1.451; *p* = 0.239). There was also no effect of genotype (F _(1, 26)_ = 1.873; *p* = 0.183) or interaction between the two factors (F _(1,26)_ = 0.318; *p* = 0.578).

#### Switching task

Mice were given training sessions where the original lever no longer delivered the outcome whilst a new lever delivered an outcome when pressed (Figure 3f). Over 4 training sessions, both groups of mice were able to switch their preference to the new active lever and reduce responding on the inactive lever (F _(1, 26)_ = 59.747; *p* < 0.001). This difference in responding increased over sessions (F _(3, 78)_ = 53.306; *p* < 0.001), with no main effect of genotype (F _(1, 26)_ = 0.027; *p* = 0.871) or any interactions with genotype as a factor (All Fs < 0.169; *p* >0.684).

To assess whether mice updated the action-outcome relationship, they were given an outcome devaluation test with access to both levers. For the inactive lever (i.e., the original lever that no longer delivered outcomes, Figure 3g), there was no effect of lever (F _(1, 26)_ = 2.620; *p* = 0.118), genotype (F _(1, 26)_ = 0.025; *p* = 0.876) or interaction between the two factors (F _(1, 26)_ = 0.606; *p =* 0.443). For the now active lever (i.e., the new reinforced lever, Figure 3h), both groups of mice were able to show outcome devaluation (valued > devalued; F _(1, 26)_ = 5.843; *p* = 0.023), with no effect of genotype (F (1, 26) = 0.024; *p* = 0.877) or interaction between the two factors (F _(1, 26)_ = 0.088; *p =* 0.769).

### Experiment 3: neuron density

The density of neurons within cortico-striatal circuits were quantified using immunohistochemistry for NeuN. There was no difference in neuron density between WT group and KD group for any of the regions examined as shown in Figure 4. This was confirmed statistically for each region within the prefrontal cortex (prelimbic, F _(1, 30)_ = 2.346; *p* = 0.136; infralimbic F _(1, 30)_ = 0.672; *p* = 0.419; lateral orbital, F _(1, 30)_ = 0.372; *p* = 0.546; anterior insular, F _(1, 30)_ = 0.350; *p* = 0.559; Figure 4a) as well as the striatum (DMS, F _(1, 30)_ = 1.484; *p* = 0.233; DLS, F _(1, 30)_ = 0.910; *p* = 0.348; NA-shell, F _(1, 30)_ = 0.755; *p* = 0.392; NA-core, F _(1, 30)_ = 0.805; *p* = 0.377; Figure 4b), and limbic regions (Figure 4c) including the ventral pallidum (F _(1, 30)_ = 0.673; *p* = 0.419), basolateral and central amygdala(F _(1, 30)_ = 0.528; *p* = 0.473 and F _(1, 26)_ = 0.292; *p* = 0.593), lateral hypothalamus (F _(1, 30)_ = 1.605; *p* = 0..215), ventral tegmental area (F _(1, 30)_ = 2.615; *p* = 0.116) and substantia nigra compacta (F _(1, 30)_ = 1.097; *p* = 0.303).

**Figure 4.**
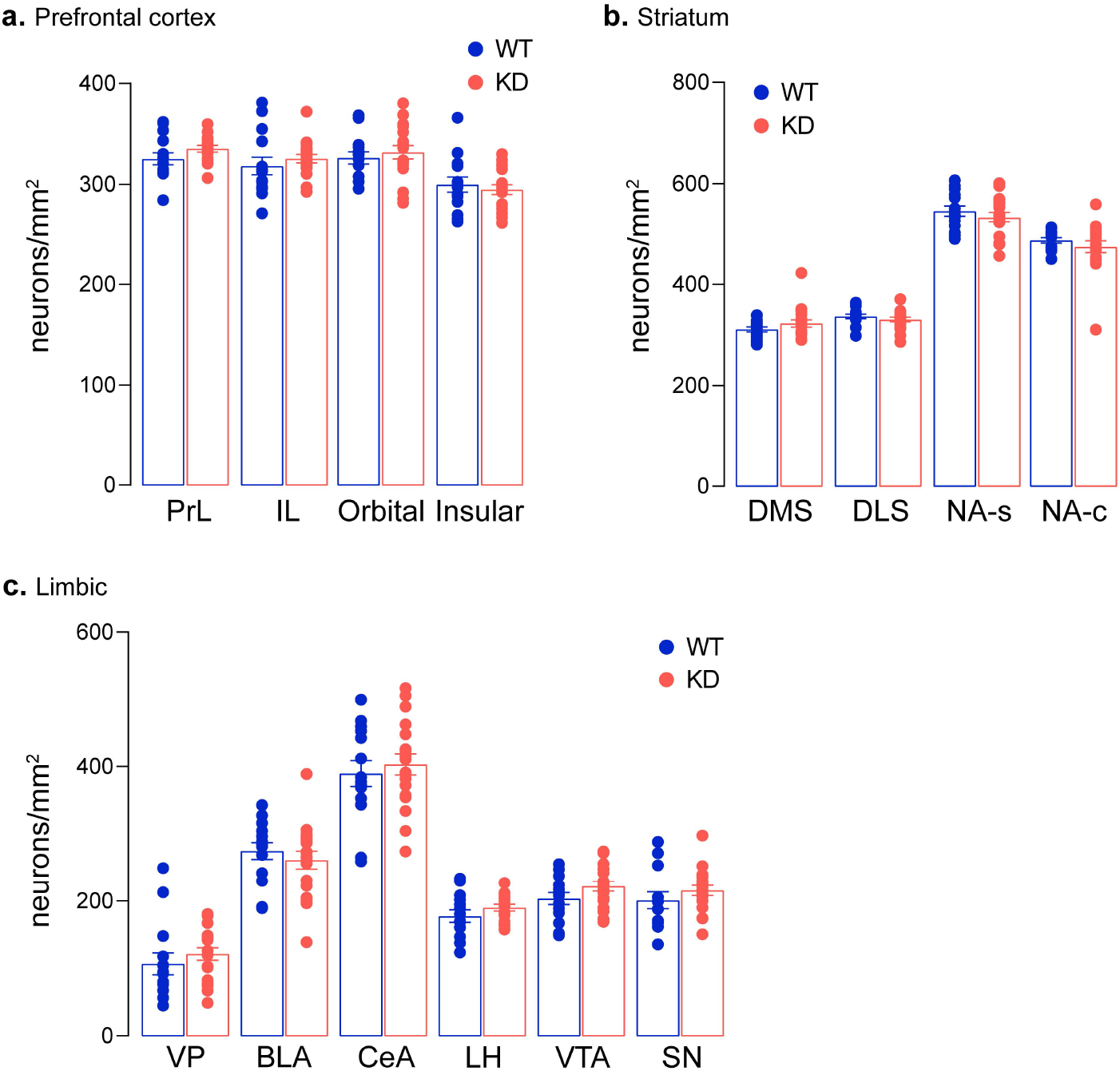
Immp2l knockdown does not alter neuronal density in cortical, striatal, or limbic regions. Neuronal density was assessed in multiple cortical, striatal, and limbic regions between WT and KD mice and no differences in neuronal density was observed. Abbreviations: Prelimbic, PrL; infralimbic, IL; orbital cortex; Orbital; insular cortex, Insular; dorsal medial striatum, DMS; dorsal lateral striatum, DLS; nucleus accumbens shell, NA-s; nucleus accumbens core, NA-c; ventral pallidum, VP; basolateral amygdala, BLA; central nucleus amygdala, CeA; lateral hypothalamus, LH; ventral tegmental area, VTA; substantia nigra, SN. Data are shown as mean ± SEM.

## Discussion

Immp2l is a mitochondrial enzyme that was knocked down in mice (Fang et al., 2017) in order to generate a model of ASD and GTS with face validity as *Immp2l* genetic alterations are often associated with ASD and GTS. The current study examined the impact of *Immp2l* knockdown (KD) on cortico-striatal structure and function. It was found that learning of neither goal-directed nor habitual instrumental actions to obtain a food outcome were altered for *Immp2l* KD mice compared to WT mice. Specifically, both KD and WT groups were goal directed in their actions following moderate instrumental training as lever preference was determined by the value of the food outcome, i.e., both groups selected the lever associated with the still valued outcome more often that the lever associated with the devalued outcome (Experiment 1). Moreover, both groups of mice were flexible in their behaviour and were able to update response-outcome contingencies following reversal learning and contingency degradation (Experiment 1), and to switch response-outcome contingencies even after prolonged training (Experiment 2). That is, lever preference was consistent with the new reversed/switched response-outcome contingencies rather than the initially learned contingencies, and with the lever where the response-outcome association was intact rather than degraded by non-contingent delivery of the pellet outcome. There was also no change in the development of habitual behaviour with both groups of mice demonstrating insensitivity to outcome devaluation after extended instrumental training (Experiment 2). However, whilst both groups of mice displayed specific PIT, there was an increased capacity for the auditory stimuli previously paired with food to promote instrumental responding for food, as KD mice increased lever responses during both the same and different CSs compared to WT mice (Experiment 1). Finally, numbers of neurons were not changed for any of the brain regions examined across the prefrontal cortex, striatum or other limbic regions, including amygdala, lateral hypothalamus or ventral tegmental area (Experiment 3). Therefore, the current study suggests that Immp2l does not alter neuron density or mediate learning that is determined by response-outcome or stimulus-response associations, but that it may play a role in sensitivity to external stimuli that drive behaviour.

### Impact of Immp2l knockdown on goal-directed and habitual behaviour

Goal-directed and habitual actions are mediated by separate neurocircuits that centre on dorsolateral striatum (DMS; caudate) and nucleus accumbens (NA), and dorsolateral striatum (DLS; putamen), respectively (Balleine & O’Doherty, 2010; Simmler & Ozawa, 2019). In rodents, these circuits have largely been determined by examining sensitivity to outcome devaluation following lesions and inactivation of specific brain regions (Balleine & O’Doherty, 2010; Lerner, 2020; Ostlund & Balleine, 2008b; Simmler & Ozawa, 2019). Several cortical and limbic structures have been determined to mediate goal-directed learning, i.e, loss of function of these regions leads to a reduction in sensitivity to outcome devaluation after moderate training (as examined in Experiment 1). These regions include prelimbic, insular, and orbital cortices, and basolateral amygdala (Bradfield et al., 2015; Corbit & Balleine, 2005; Corbit et al., 2013; Hart et al., 2018; Ostlund & Balleine, 2008a; Parkes et al., 2015). Unique cortical and limbic structures have also been determined to mediate habitual actions where loss of function in these structures reduces the insensitivity to outcome devaluation that occurs for well-learned actions after extended training (as examined in Experiment 2). This includes infralimbic cortex, dopaminergic nigral neurons, central amygdala, and lateral hypothalamus (Becchi et al., 2021; Bingul et al., 2022; Coutureau & Killcross, 2003; Lingawi & Balleine, 2012; Marquez et al., 2020; Yin et al., 2004). In the current study, there was no change to numbers of neurons in these brain structures, and no change to acquisition or expression of goal-directed or habitual learning following knockdown of *Immp2l*.

It has been proposed that dysregulation in these neurocircuits may underlie some of the symptoms of ASD and GTS including tics, higher-order repetitive behaviours and compulsive behaviours (Alvares et al., 2016; Delorme et al., 2016; Gillan et al., 2011; Hadjas et al., 2019; Scholl et al., 2022; Singer, 2013). Indeed, both ASD and GTS are characterised by neuroanatomical changes in these circuits post-mortem, including reduced neuron density and numbers of neurons in striatum (Kalanithi et al., 2005; Kataoka et al., 2010; Wegiel et al., 2014) and increased numbers of neurons in prefrontal cortex in ASD (Courchesne et al., 2011; Falcone et al., 2021). Furthermore, changes in size of the caudate (i.e., DMS) correlate with severity of tics, repetitive behaviours and compulsions (Bloch et al., 2005; Langen et al., 2009). In addition, when tested using translation versions of the procedures used in the current study, individuals with ASD displayed reduced goal-directed learning as their actions were not determined by outcome value, and individuals with GTS showed enhanced habits and an over-reliance on stimulus-response associations (Alvares et al., 2016; Scholl et al., 2022). Thus, the current study suggests that immp2l is not involved in these aspects of ASD or GTS.

Of note, potentiated habitual behaviour has been demonstrated in a genetic mouse model of OCD (Sapap3 KD), and thus habitual actions may model repetitive actions that do not have well-defined goals (i.e., compulsive repetitive behaviour in OCD)(Hadjas et al., 2019). Increased self-grooming behaviour was also seen with these mice, which is used as a measure of lower-order repetitive or compulsive behaviour (Crawley, 2007; Hadjas et al., 2019; Jiang & Ehlers, 2013). Thus, there may be a possible link between lower and higher order repetitive behaviour. We have previously shown that lower-order repetitive behaviour was not altered in *Immp2l* KD mice, as there was no effect on self-grooming behaviour or marble burying behaviour (Kreilaus et al., 2019; Lawther et al., 2022). Thus, it would be of interest to determine whether increased self-grooming behaviour predicts altered habitual behaviour in other mouse models of ASD and GTS (e.g., Cntnap2- and Shank-targeted mice) (Fuccillo, 2016; Peca et al., 2011; Penagarikano et al., 2011), and thus may be dependent on the same neurocircuitry. Further, there are ASD and GTS mouse models with recognised neuroanatomical and functional changes in the striatum, (again both Cntnap2- and Shank-targeted mice among others)(Fuccillo, 2016; Isaacs & Riordan, 2020; Peca et al., 2011; Penagarikano et al., 2011), which are thus excellent candidates for testing the behavioural procedures used in the current study to attempt to model the imbalances in goal-directed and habitual behaviour seen in ASD and GTS.

### Impact of Immp2l knockdown on new learning and behavioural flexibility

*Immp2l* KD also did not impact reversal learning, contingency degradation, switching, or extinction learning in the current study. These procedures require cognitive flexibility where there must be inhibition of the initially learned response-outcome association, and then new learning of an updated response-outcome relationship (Bradfield et al., 2013; Furlong et al., 2016; Gourley et al., 2010; Ostlund & Balleine, 2008b). Deficits revealed by these procedures are indicative of cognitive rigidity and correlate with repetitive behaviour in individuals with ASD, and are thus used in rodents to model higher-order repetitive behaviours including insistence in sameness, resistance to change, rigid behaviour, and obsessive behaviour (Alvares et al., 2016; Crawley, 2007; D’Cruz et al., 2013; Lewis et al., 2007; Tanimura, Yang, & Lewis, 2008; Whitehouse, Curry-Pochy, Shafer, Rudy, & Lewis, 2017). The neural structures associated with reversal learning and contingency degradation include the orbital and prelimbic cortices, dorsomedial striatum and basolateral amygdala (Balleine & Dickinson, 1998; Bradfield & Balleine, 2017; Bradfield et al., 2013; Fresno, Parkes, Faugere, Coutureau, & Wolff, 2019; Ostlund & Balleine, 2008a), where neuron density was also not changed in the current study. Furthermore, acquisition of instrumental responses (during training) was also not changed in the current study, which is used to model cognitive disability (Crawley, 2007) and is co-morbid with ASD (Giovinazzo et al., 2013; Larson et al., 2010). Thus, the current study suggests that Immp2l is not involved in initial learning or updating learning, which have been proposed to contribute to higher-order repetitive behaviours (Alvares et al., 2016; Crawley, 2007; D’Cruz et al., 2013; Lewis et al., 2007; Tanimura et al., 2008; Whitehouse et al., 2017).

### Impact of Immp2l knockdown on Pavlovian stimuli

Whilst the current study suggests that Immp2l is not involved in instrumental learning, it was demonstrated that the expression of instrumental behaviour can be invigorated by Pavlovian stimuli when *Immp2l* was knocked down. This increase in responding during PIT was general rather than specific to learned associations, as it occurred for a CS paired with a different outcome (than that of the instrumental behaviour) in addition to the CS paired with the same outcome (as the instrumental behaviour). The observed general increase in responding to the CSs was not likely to be due to changes in motivation for food as *Immp2l* KD mice did not demonstrate altered responses at any other times, i.e., when food was delivered during training (Pavlovian or instrumental training) or when food was not delivered during extinction (prior to PIT). There was also no difference in consumption of food during the free-access devaluation periods. The general increase in responding was also not due to changes in learning as specific PIT was demonstrated by KD mice (Laurent et al., 2012; Leung & Balleine, 2015; Ostlund & Balleine, 2008b), which further indicates that there were no deficits in learning for KD mice, this time learning of Pavlovian associations, response-outcome associations, and outcome-response associations (Holmes et al., 2010). Thus, it may be that KD mice are more sensitive to Pavlovian auditory stimuli than WT mice resulting in increased capacity for environmental cues to control behaviour.

Individuals with ASD and GTS often demonstrate altered sensory processing and sensory hypersensitivity, including to auditory stimuli (Beste & Munchau, 2018; Isaacs & Riordan, 2020; Kargas, Lopez, Reddy, & Morris, 2015; Schulz & Stevenson, 2019). It has been proposed that tics may occur because of increased drive from internal stimuli (e.g., premonitory urges) and from external stimuli, which can result in ‘perception-action’ associations and ‘stimulus-induced tics’ (Beste & Munchau, 2018; Cavanna et al., 2020; Isaacs & Riordan, 2020). Similarly, environmental stimuli can influence the occurrence of repetitive behaviour, and sensory hypersensitivity predicts repetitive behaviour in ASD and GTS (Kargas et al., 2015; Schulz & Stevenson, 2019) and insistence in sameness in ASD (Black et al., 2017). Altered responses to auditory stimuli are typically measured via acoustic startle, sensory gating, and responsiveness to sensory cues as is measured in PIT (Crawley, 2007; Isaacs & Riordan, 2020). We have previously shown that *Immp2l* KD does not change acoustic startle or pre-pulse inhibition in male mice (Kreilaus et al., 2019) which suggests that these mice are not more sensitive to (unconditioned) auditory stimuli generally, and that Pavlovian auditory stimuli may uniquely influence behaviour.

In any case, PIT is known to be dependent on NA and associated neurocircuits including ventral tegmental area (VTA), ventral pallidum, basolateral amygdala and prefrontal cortex, with these structures playing different roles in different aspects of PIT (Corbit & Balleine, 2005; Holmes et al., 2010; Keistler, Barker, & Taylor, 2015; Laurent et al., 2012; Leung & Balleine, 2015; Lex & Hauber, 2008; Ostlund & Balleine, 2008a; Saddoris, Stamatakis, & Carelli, 2011). Of particular interest, disrupting communication between the caudal VTA and ventral pallidum increase lever responses during the different CS and pre-CS periods, as occurred in the current study (but also reduces specific PIT)(Leung & Balleine, 2015). Further, administration of cocaine (an indirect dopamine agonist) to rodents produces a similar profile of PIT to that of the current study where specific PIT was maintained along with increased responding during CSs and pre-CS periods (Saddoris et al., 2011). Conversely, inactivation of the VTA (and presumably dopamine neurons) reduces responding during both pre-CS and CS periods (Corbit, Janak, & Balleine, 2007). Hence, Immp2l-induced changes to PIT may be mediated by dopamine, which is produced in VTA and has been shown to act at the NA to influence PIT (Lex & Hauber, 2008; Saddoris et al., 2011). Thus, the mesolimbic circuit is a prime candidate for dysfunction following *Immp2l* KD. In accordance, we have shown prior that *Immp2l* KD mice demonstrate increased locomotor responses to d-amphetamine (also an indirect dopamine agonist)(Kreilaus et al., 2019) which is also a mesolimbic-mediated behaviour. Of note, lesions of NA increase locomotor responses to d-amphetamine, and infusions of d-amphetamine into the NA (but not dorsal striatum) increases locomotion (Parkinson et al., 1999; Staton & Solomon, 1984) as well as cue-driven lever responses (Wyvell & Berridge, 2000), which further suggest that the NA is a likely site of actions for our effects.

Whilst neurons were not altered in any of the regions examined, including NA and VTA, it may be that neurons were in a pre-degenerative dysfunctional state not detected by NeuN immunostaining given that a prior study using a different *Immp2l* KD mouse model showed that neuron density (in the cerebellum) was only reduced in older and not younger mice (i.e., older than 20 months)(Liu, Li, & Lu, 2016). Alternatively, *Immp2l* KD has also been shown to increase oxidative stress (Lu et al., 2008) and dysregulate genes involved in central nervous system development and cell proliferation (Gokoolparsadh et al., 2017), which may lead to dysfunction in these brain regions and subsequent behavioural changes. Other potential neural changes that should be examined in future studies are those related to neuroinflammation (e.g., increased microglia) and alterations to the dopamine system (e.g., increased striatal dopamine-2 receptors), as both have been shown to be altered in the brains of individuals with ASD or GTS (Beste & Munchau, 2018; Brandenburg et al., 2020; Hienert, Gryglewski, Stamenkovic, Kasper, & Lanzenberger, 2018; Ueda & Black, 2021; Varghese et al., 2017).

## Conclusion

In summary, the current study showed that *Immp2l* knockdown does not alter goal-directed or habitual learning, or the ability to learn new rules associated with actions, but that Pavlovian stimuli can invigorate instrumental responding and thus there may be increased capacity for environmental cues to control behaviour. This increase in stimulus-controlled behaviour may contribute to tics and repetitive behaviour, seen in ASD and GTS, which are proposed to be, in part, stimulus-driven behaviours (Cavanna et al., 2020; Delorme et al., 2016; Isaacs & Riordan, 2020; Kargas et al., 2015; Schulz & Stevenson, 2019). There were no alterations to numbers of neurons in any of the cortical, striatal or limbic regions examined, including regions where Immp2l is expressed at high levels, i.e., striatum, hypothalamus, amygdala, and substantia nigra (Bertelsen et al., 2014). Hence, other changes are likely to have occurred in the brain, especially in regions that regulate PIT such as VTA-NA mesolimbic circuit, which should be a key focus for future studies examining neural changes that occur following *Immp2l* KD. It should also be noted that the current study was restricted to male mice given sexual dimorphism of ASD and GTS (Burd et al., 2009; Freeman et al., 2000; Kadesjo & Gillberg, 2000), and it may be that *Immp2l* KD in female mice would lead to different outcomes than those of male mice in the current study (as occurred in Kreilaus et al., 2019 where sensitivity to d-amphetamine was not seen in female mice unlike male mice). Regardless, our mouse model, with face validity, suggests that knockdown of *Immp2l* alone does not contribute to an imbalance of goal-directed versus habitual behaviour or associated neurocircuits seen in individuals with ASD and GTS. However, whilst *Immp2l* KD in isolation does not alter these phenotypes, it may be that it increases susceptibility so that deficits emerge if *Immp2l* KD occurs in combination with other known genetic or environmental risk factors (Lewis et al., 2007; Moy & Nadler, 2008; Ueda & Black, 2021; Varghese et al., 2017). Given the complexity of genetic and environmental factors that contribute to ASD and GTS, mice with specific gene knockdown are an excellent tool for linking specific genes to functional outcomes (Moy & Nadler, 2008). To our knowledge this is the first time that procedures examining sensitivity to outcome devaluation and PIT have been applied to a mouse model of ASD or GTS. Thus, the current study highlights the utility of applying these procedures to other mouse models of ASD and GTS to provide insight into the processes that underlie the development of higher-order repetitive behaviours and tics.

## Acknowledge

We wish to acknowledge Keira McLoskey and Saoirse Patton for their contribution to the laboratory work.

## Notes

**Grants:** This work was supported by a Strategic Initiative Project from the Australian Research Council (CE140100007) awarded to TMF, SM, BWB and GP, a Sydney Partnership for Health Education Research and Enterprise (SPHERE) Grant awarded to TMF, AKW, VE, BWB, RC, SM, and BKL, and AW08 Ingham Institute Mental Health Research Award 2018 to RC and VE.

### Competing Interest Statement

The authors have declared no competing interest.

